# Street light choice matters: impacts of presence and color on wild plants

**DOI:** 10.64898/2026.02.16.703286

**Authors:** E. Castrop, P. M. van Bodegom, E. F. Strange, S. H. Barmentlo

## Abstract

Anthropogenic light at night (ALAN) is rapidly expanding, affecting wildlife and ecosystems. Color is an important factor in designing responsible outdoor lighting strategies, yet color choice effects on plants remain largely unknown. We investigate how different colors of street lighting influence the morphology and nighttime assimilation rates of two wild plant species, *Hypochaeris radicata* and *Rumex acetosa*. Using a semi-natural mesocosm setting, plants were exposed to nightly red, green, and white streetlights, with a dark control, over an eight-week period. Red light significantly increased leaf length and petiole elongation compared to other treatments. Nighttime transpiration and stomatal conductance also strongly increased under both red and white light. These findings highlight that ALAN, particularly red and white light, can disrupt plant growth and physiological processes, which can potentially affect overall ecosystem dynamics. There’s also a species-specific effect to morphological parameters which is absent for the assimilation parameters. Striking a balance between functionality and ecological impact is challenging, given that certain light colors negatively affect plants, while those same colors may be preferred by other organisms. A deeper understanding is needed to guide informed decisions on plant and wildlife protection.

## 1. Introduction

Anthropogenic light at night (ALAN) is growing faster than many other key environmental pressures caused by human activities and covers areas just as extensive (Gaston et al., 2021). Global light pollution has been estimated to increase by 2.2% per year for the last decade with no expectation of slowing (Kyba et al., 2017; Sánchez De Miguel et al., 2021). Streetlights are recognized as a major contributor to light pollution, accounting for an estimated 20–40% of its overall impact (Kyba et al., 2021). ALAN can affect organisms and interactions between them, cause habitat loss and negatively impact foodwebs (Jägerbrand & Spoelstra, 2023, Bennie et al., 2016; Bucher et al., 2023; Singhal et al., 2018; Heinen, 2021, Gaston et al., 2021). Although ALAN has been shown to affect animals and their interactions, plants have received less attention, particularly regarding light color. Understanding plant responses is essential, as ALAN has the potential to disrupt growth and reproductive patterns, interfere with seasonal rhythms, and alter physiological processes such as stomatal dynamics and respiration (Bennie et al., 2018; Johnston et al., 2025; Kwak et al., 2018). ALAN has also been linked to decreased plant growth and higher water demand, connecting light pollution to other environmental stressors (Hey et al., 2020). The broader effects of ALAN, especially across different wavelengths, on plant morphology and physiology are still not fully understood (Bennie et al., 2015; Heinen, 2021). Therefore, developing holistic mitigation strategies requires knowledge of impacts across all trophic levels, including plant responses.

Existing outdoor lighting guidelines include color-related recommendations, such as those proposed by the Dark Sky Association within its five principles of responsible outdoor lighting (Schaefer, 2020). Also, red light has received attention as a potentially ecologically favorable lighting color (Durmus et al., 2024). But this is primarily based on animal studies, not plants. . As light sources increasingly transition to solid-state light emitting diode (LED) technology, characterized by broader spectral outputs, this shift potentially provokes differing responses in plant biota. Plants exhibit specific responses depending on the color spectrum where red-blue illumination notably enhances photosynthetic rates, stomatal conductance, and consequently, plant development and yield (Camejo et al., 2020; Singhal et al., 2018). The spectral composition of daytime light strongly affects the efficiency of light absorption and CO_2_ fixation and is an important determinant of growth (Lazzarin et al., 2021). During daytime in natural canopies, vegetation absorbs red and blue light for photosynthesis and reflects far-red light. Underneath the canopy, the resulting reduced red and blue color is perceived by photoreceptors and signals shading and light competition from neighboring plants. Plants respond to a low red: far-red ratio with an increased growth response (e.g. elongation and increased leaf area) to avoid the shade (Meijer et al., 2022). Similar processes may occur under low light intensities typical of ALAN. However, these responses have not yet been studied in semi-natural field settings. Therefore, we aimed to assess if the morphology and assimilation of wild plant species are impacted by different colors of light emitted by streetlights at night. We placed two wild plant species, *Hypochaeris radicata* and *Rumex acetosa*, within a semi-natural mesocosm setting underneath different colored streetlights in dark nature reserves for 8 weeks. Based on current knowledge, we expect that plants exposed to red or white light at night have an increased growth and nighttime transpiration compared to green light and a dark control. This study will give decision-makers insight into how ALAN, and the choice of light color, can affect plant functioning of urban and near-urban ecosystems.

## 2. Methods

### 2.1 Study site

The study was conducted at two of the experimental sites previously initiated by Spoelstra et al., 2015; Radio Kootwijk (52°11’20’’N 5°48’17’’E) and Lebret’s Hoeve (52°5’53’’N 5°55’59’’E) in The Netherlands. Subsequently referred to as locations one and two respectively. In selecting these sites, the absence of other sources of anthropogenic light (van Langevelde et al., 2011) was a requirement. The experimental field sites were in forest-edge habitats, which included both forested and open-field areas. At each site, there were 4-meter-tall lampposts arranged in four 100-meter rows, positioned perpendicular to the forest edge. Each row contained two lampposts in the open field, one at the forest edge, and two within. For this experiment, only lampposts situated in the open-field areas were used, as these minimize potentially disrupting shadow effects. Each row was randomly assigned a color (green, red or white), with one row left unlit as a control. All colors emitted 8.2 lux (SE = 0.3) at ground level, consistent with typical street lighting in The Netherlands.

During the experimental period from April till the end of June 2023, the average monthly rainfall was 50 mm. Daily temperatures averaged 13 °C, with about 8 hours of sunshine per day. This data was sourced from the Deelen weather station (coordinates 52°3’18”N 5°52’19.2”E), operated by the Royal Netherlands Meteorological Institute (KNMI, 2022).

### 2.2 Plant species

In this study, we worked with two plant species: *H. radicata* and *R. acetosa*. These two species were chosen due to their natural occurrence at the test sites, their low palatability to herbivores, and the requirement that the leaves of the plants needed to be at least 4 cm^2^ in size to facilitate all measurements (see section 2.3). The plants were purchased as seedlings from a nursery specialized in wild plants (Cruydthoeck, n.d.). Seedlings were repotted into pots measuring 12 cm in diameter and 9 cm in height and were then left for a two-week acclimatization period in an open greenhouse at the Netherlands Institute of Ecology (NIOO), Wageningen. After two weeks, all plants had survived the acclimatization period. For each treatment, 12 plants were randomly selected and placed into a box measuring 800 x 600 x 200 mm. Each plant was placed in an individual pot within the box. The box itself was then filled with white sand for stability and soil temperature control.

### 2.3 Experimental setup

The experiment was designed to test the effects of different colors: red, white, green (see Figure s1 in Spoelstra et al,. 2015 for the full spectral composition) and a dark control. At both field sites, one lamppost of each of the four treatments was located. A box containing 12 plants (replicates) was placed directly beneath each lamppost. We placed a higher number of plants than we actually measured to account for potential losses by large herbivores. Throughout the experiment, regular watering ensured that water availability was not a limiting factor. The duration of the experiment spanned two months (including the acclimation time, see section 2.2).

Morphological characteristics were measured weekly in 5 randomly selected plants per species per box (for a total of 10 measurements per treatment since we used 2 locations). Because of time constraints during the night, we did not measure the full 12 plants per box. For simplicity, we only present the morphological data at the end of the experiment since effects of time were non-apparent (i.e. after 29 days of exposure, all time series data are provided in the data repository, Castrop et al., 2025). Measurements were conducted using a caliper: leaf width was recorded at the widest point, leaf length from base to apex, and petiole length (measured only in *R. acetosa*, as it is absent in *H. radicata*) from the soil to the leaf base. Leaf herbivory was assessed by counting the number of leaves with feeding scars per plant and estimating the percentage of the total leaf surface area eaten.

Gas flux rates, including transpiration rates (mmol H_2_O m^−2^ s^−1^) and photosynthetic rates (μmol CO_2_ m^−2^ s^−1^), were measured in 4 randomly selected plants per box (for a total of 8 measurements per treatment per sampling moment) during both day and night periods. Daytime measurements were taken at noon, while nighttime measurements were taken one hour after astronomical sunset. For both plants we selected fully grown leaves that were undamaged by any means. For *H. radicata* we selected the second youngest leaf that was fully grown at the moment of the first sampling. For *R. acetosa*, we selected fully grown leaves comparable in size between plants. All selected leaves were labeled to make sure subsequent measurements were performed on the same leaf. We placed a TARGAS-1 Portable Photosynthesis System by PP Systems leaf chamber (4.5 cm^2^) over the selected leaves so the chamber was completely filled. The chamber was left on the leaf for 5 minutes for stabilization after which the measurement was taken. Due to weather conditions, access constraints of the nature reserve and availability of the equipment, this process was performed one week apart for the different locations during the last two weeks of the experiment (i.e. after 22 and 29 days of exposure respectively).

### 2.4 Statistical analyses

Statistical analyses were performed using R-Studio (version 4.3.2). We applied mixed linear models to examine the variables leaf length, leaf width, petiole length, herbivory, transpiration, photosynthetic rate, and stomatal conductance. In each model, species and color were treated as fixed factors, including their respective interactions while location was a random factor. If significant interactions were detected, analyses were conducted separately for each plant species. Post hoc Dunnett’s tests were used to compare the effects of color against the dark control. Results are presented with error bars representing the standard error of the mean (± SE), and statistical significance was determined with a threshold of alpha = 0.05.

## 3. Results

### 3.1 Morphological changes

The analysis of morphological changes revealed the following results: The light treatments strongly affected leaf length (figure 1a) (P < 0.001). Plants had longer leaves under red light than at the dark control (by 11% for *H. radicata* and 8% for *R. acetosa*, P = 0.05), while there was no trend visible for the other comparisons, including control vs. green, and control vs. white. For petiole length (figure 1c), *R. acetosa* plants under red lights showed a significantly higher petiole length than at the dark control (an increase of 34%, P < 0.01), with no significant differences found in comparisons between control and green light or control and white light. Additionally, leaf width and leaf area (figure 1b and d) showed no significant effects of different light colors. Overall, leaf herbivory was relatively low with a maximum of 5% and 33% of *H. radicata* and *R. acetosa* individuals only slightly affected at any given time. Of those individuals, the total leaf area affected was on average 11%. We observed no significant effect of light color on the number of plants that were herbivorized, the number of leaves affected, and the total area affected (p > 0.05 for all comparisons), which is why we excluded these parameters from further mention.

**Figure 1.**
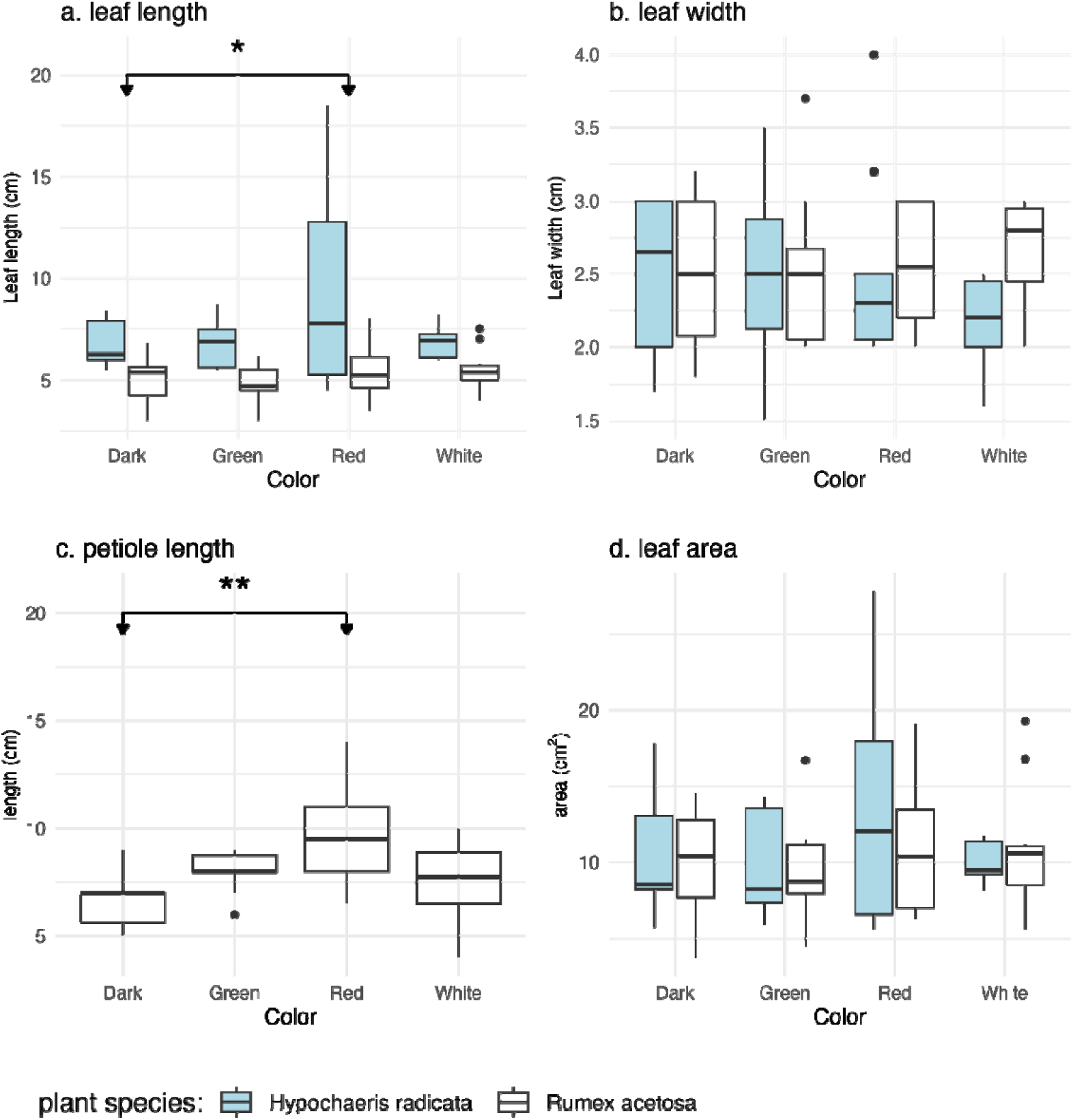
Boxplots illustrating the effects of light color (green, red and white versus dark) on various leaf traits. Figure A displays leaf length (in cm, n=10) distributions during the day, Figure B presents leaf width (in cm, n=10), Figure C shows petiole length (cm, n=10) distributions, and Figure D showcases leaf area (cm^2^, n=10) distributions after 29 days of exposure. In each figure, blue-colored boxplots represent *H. radicata*, while white boxplots represent *R. acetosa*.

### 3.2 Assimilation at night

There were significant differences in the effects of various light conditions on the transpiration and assimilation. For transpiration at night (figure 2a), the comparison between the dark control and red light revealed an increase in response under red light (an increase of 104%, P < 0.01). Transpiration at night was higher at white light conditions than at the dark control (an increase of 57%, P = 0.01) and when white and green light conditions are compared (an increase of 122%, P < 0.01). The comparisons between red and white light, as well as between dark and green light, did not show significant differences. Night stomatal conductance was strongly affected by the light treatments (figure 2d) (P < 0.01) and the results revealed significant differences in the effects of the red-light conditions compared to the dark control. The stomatal conductance at the dark control was significantly lower than under red light (100%, P < 0.01), while green light (SE = 7.74, P = 0.13) and white light (SE = 7.74, P = 0.22) did not lead to changes in stomatal conductance. The results for photosynthetic rate (figure 2f) indicate no significant differences in the effects of any of the light conditions compared to a dark control.

**Figure 2.**
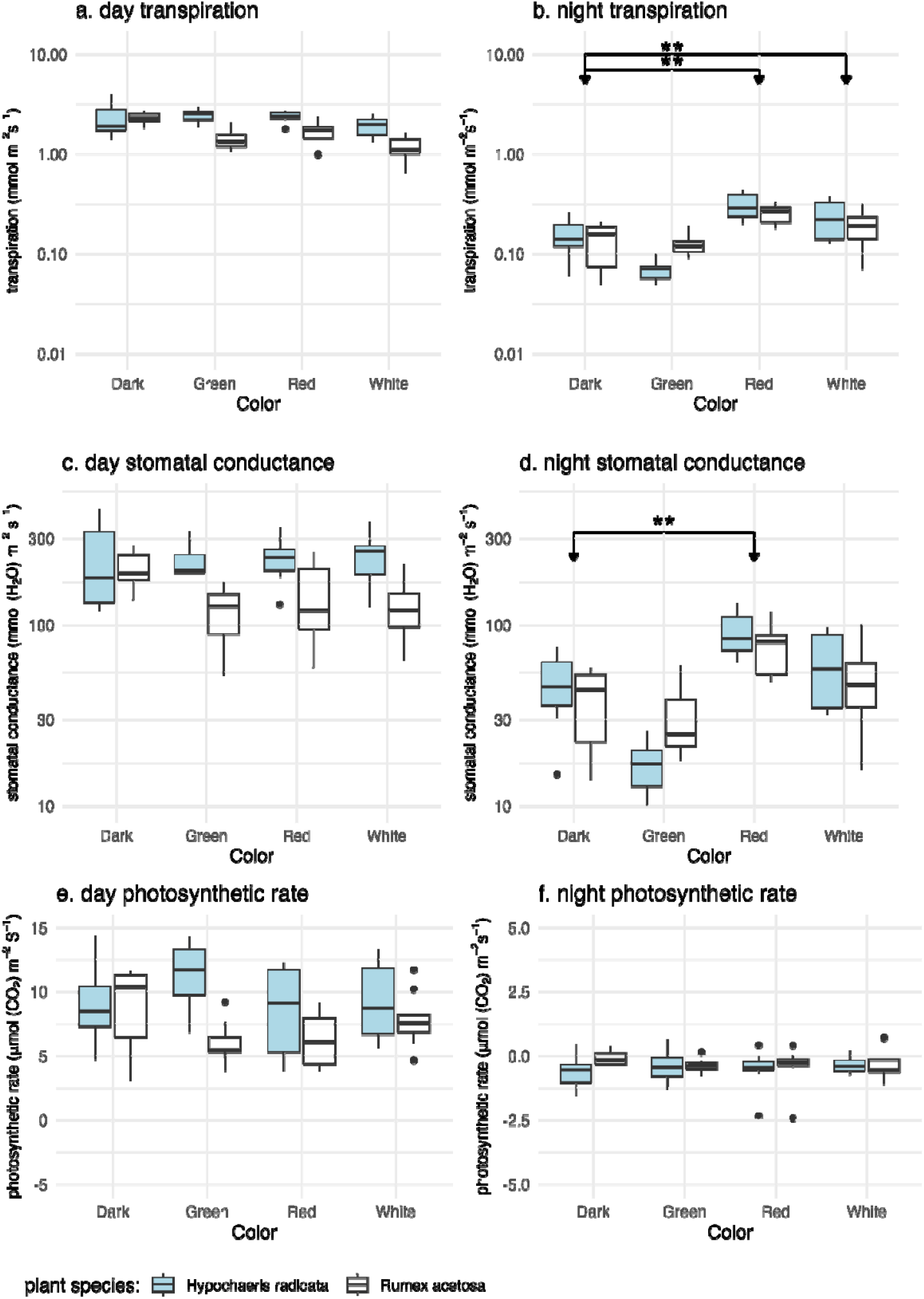
Boxplots of effects of light color (green, red and white versus dark) on transpiration rates (mmol (H□O) m^−2^ s^−1^) during the day and night (n=8, Figures A and B), stomatal conductance (mmol (H□O) m^−2^ s^−1^) during the day and night (n=8, Figures C and D), and photosynthetic rates (μmol (CO□) m^−2^ s^−1^) during the day and night (n=8, Figures E and F) after on average 26 days of exposure. Blue-colored boxplots represent *H. radicata*, while white boxplots represent *R. acetosa*.

### 3.3 Assimilation at daytime

The results for the daytime transpiration and photosynthetic rate (Table 1) show a significant interaction between species and color: regarding transpiration data (Figure 2a), we see that for *H. radicata*, the output is unaffected by color. However, for *R. acetosa*, we observed a significant decrease in transpiration under all light conditions compared to the dark control: a 36% decrease for green light, a 27% decrease for red light, and a 50% decrease for white light (SE= 11.48, P < 0.01). In the photosynthetic rate data (Figure 2e), a significant interaction between species and light color was observed. Under red and white light, no differences were detected between species or compared to the dark control. However, under green light, we observed a 28% increase in *H. radicata* and a 32% decrease in *R. acetosa* compared to the dark control (SE=9.55, P = 0.02). Daytime stomatal conductance (figure 2c) did not show significant differences when compared the dark control to the colored light conditions.

**Table 1.** The statistical test outcomes for the comparison of morphological and assimilation parameters across plant species and color treatments.

|  | <u>color</u> |  |  | <u>Species</u> |  |  | <u>interaction</u> |  |  |
| --- | --- | --- | --- | --- | --- | --- | --- | --- | --- |
| <u>morphology</u> | DF<br>(residuals) | Chisq | P<br>value | DF<br>(residuals) | Chisq | P value | DF<br>(residuals) | Chisq | P value |
| *Leaf length* | 3 | 14.38 | <0.001 | 1 | 4.84 | 0.028 | 3 | 5.74 | 0.125 |
| *Leaf width* | 3 | 4.04 | 0.257 | 1 | <0.01 | 0.880 | 3 | 4.00 | 0.262 |
| *Petiole length\** | 3 | 18.99 | <0.001 | NA | NA | NA | NA | NA | NA |
| *Leaf area* | 3 | 6.10 | 0.107 | 1 | 0.14 | 0.7096 | 3 | 2.26 | 0.521 |
| <u>Assimilation</u> |  |  |  |  |  |  |  |  |  |
| **Night** |  |  |  |  |  |  |  |  |  |
| *Transpiration* | 3 | 51.77 | <0.001 | 1 | 0.90 | 0.342 | 3 | 6.10 | 0.107 |
| *Photosynthetic rate* | 3 | 6.11 | 0.106 | 1 | 1.29 | 0.256 | 3 | 3.63 | 0.305 |
| *Stomatal conductance* | 3 | 48.43 | <0.001 | 1 | 1.73 | 0.189 | 3 | 5.06 | 0.167 |
| **Day** |  |  |  |  |  |  |  |  |  |
| *Transpiration* | 3 | 7.42 | 0.060 | 1 | <0.01 | 0.905 | 3 | 11.48 | <0.001 |
| *Photosynthetic rate* | 3 | 6.66 | 0.084 | 1 | <0.01 | 0.868 | 3 | 9.55 | 0.023 |
| *Stomatal conductance* | 3 | 1.79 | 0.617 | 1 | 0.69 | 0.408 | 3 | 3.62 | 0.306 |

\* Only tested for *R. acetosa*, because of absence of a clear petiole in *H. radicata*.

## 4. Discussion

We investigated the effects of different streetlight colors on day- and nighttime assimilation rates and morphological changes in two plant species in a semi-natural setting. We found streetlight color choice matters for plant performance with a significant increase in leaf length under exposure to red light for both species and petiole elongation in *R. acetosa*. Moreover, both red and white light triggered a doubling of nighttime transpiration and stomatal conductance in both plant species.

To explain these results, we consider the impacts of low daytime illumination, since there is very limited literature on nighttime low intensity illumination in field settings. The spectral composition of daytime light strongly affects the efficiency of light absorption and CO_2_ fixation and is an important determinant of growth (Lazzarin et al., 2021). Underneath canopies, there are low light intensities with a reduced red: far red ratio during daytime (Meijer et al., 2022; Roig-Villanova et al., 2019). The resulting reduced red: far-red ratio is perceived by photoreceptors which signal shading and light competition from neighboring plants. Plants respond to a low red: far-red ratio with an increased growth (e.g. elongation) response to avoid the shade (Roig-Villanova et al., 2019). The shade avoidance observed in plants under canopies aligns with our findings of increased leaf length and petiole length. However, it was the red-light treatment which showed the most pronounced effects, despite this treatment involving a higher red-to-far-red ratio. Our results thus show that plants are adjusting their morphology in response to different wavelengths and light intensities exhibited by streetlights at night.

The assimilation data showed significant effects under white and red-light exposure on both nighttime transpiration and stomatal conductance for both tested plant species. The red light does not include a blue component while the white light has both red and blue components. Both the blue and red parts of the spectrum play a central role in the signaling pathways associated with stomatal dynamics in response to light quantity and quality (Matthews et al., 2020; Shimazaki et al., 2007), but stomatal opening in response to light is induced by different mechanisms depending on the wavelength. Blue light is known to directly influence the guard cells, while red light requires a higher light intensity to trigger a response, likely due to the involvement of a more complex signaling pathway (Kinoshita & Shimazaki, 1999; Roelfsema et al., 2002). Our results suggest that both the blue and red photoreceptors could be triggered by ALAN coming from streetlights. Red light requires higher light intensity to achieve the same level of visibility as white light because of its lower luminous efficiency. This is due to the mismatch between red wavelengths and the human eye’s spectral sensitivity functions, both photopic (daylight) and scotopic (low-light) (Mander et al., 2023). However, we observed similar responses in plants for both the red and white light treatments, suggesting that both the red and blue components of the light played a role in the stomatal conductance response. Thus, under streetlight conditions, red light adapted to the needs of human vision is just as harmful to plants as white light.

Green light also showed a tendency for decreased stomatal conductance compared to the dark. While most research focuses on how red and blue light affect stomata and photosynthesis, green light appears to play a role as well. Green light might inhibit the opening of stomata caused by blue light (Talbott et al., 2006; Aasamaa and Aphalo, 2016), aligning with the lower stomatal conductance observed in our study. However, the literature on low-intensity green light is largely absent, so our results should be considered indicative and suggest an interesting angle for future research.

The nighttime photosynthetic rate did not display any differences between treatments. Similar results were found in *Eucalyptus camaldulensis* (Lockett et al., 2022). It was expected that photosynthetic rate responses are closely linked to the stomatal conductance (which did respond to ALAN) since the stomatal opening regulates the influx and outflux of CO_2_. The observed decoupling might have been made possible because photosynthesis and stomata do not react to changes in light intensity and quality at the same speed (Shimazaki et al., 2007). However, this mechanism would only explain differences after a short adaptation time. Since our measurements were taken after hours of darkness it is more likely that the change in stomatal conductance is independent of photosynthetic rate. In a controlled laboratory experiment, increased stomatal conductance was found to be influenced by signals unrelated to photosynthesis, varying under different lighting conditions. There was an increase in transpiration and stomatal conductance, but not in carbon accumulation (Baroli et al., 2008). This may also be occurring with light pollution. In combination, these results suggest that ALAN has a low enough intensity to trigger a stomatal response and increase transpiration, but not to induce a photosynthetic response.

The decoupling of physiological responses can negatively impact plant fitness by causing increased water loss and energy expenditure without the benefit of photosynthetic yield, potentially leading to stress and reduced growth. In turn, reduced growth would lead to a decrease in overall plant biomass. Such decreases in biomass have also been observed in other studies (Bucher et al., 2023), where they found that biomass was reduced by 33% in the highest ALAN treatment compared to the control. If the observed assimilation rates in our two species are representative for other species, then the decrease in biomass can affect entire ecosystems, as plant biomass serves as the primary energy source for the system. If primary production is affected, it can have a bottom-up effect on the ecosystem, impacting fauna as well (Heinen, 2021). Reduced plant biomass can lead to less food availability for herbivores, which in turn affects predators. Disruption of plant growth and health can alter habitats, impacting species that rely on these plants for shelter and food. It is plausible that such effects are widespread in the environment, with artificial light at night impacting considerable areas of urban, suburban, and roadside vegetation globally (Falchi et al., 2016).

Effects of ALAN on plant biomass may be highly species-specific, given that artificial light at night interacts with various parameters, affecting plant biomass in complex ways. The response of those parameters may depend on the environmental conditions. For instance, Hey et al. (2020) found increased biomass under ALAN, particularly when soil moisture was limited. Additionally, ALAN influences factors such as soil biology, herbivory, and nutrient availability (Cesarz et al., 2023; Cieraad et al., 2022), which can further impact biomass outcomes. If species respond differently to these conditions, ALAN will affect the competition among plant species, which will affect plant cover and plant composition (Liu et al., 2022; Murphy et al., 2022). These findings highlight the complex nature of ALAN’s impacts. Light pollution is rarely the only factor; climate change, excess or shortage of water, and soil pollution are often present in the same areas. Our results may help clarify some aspects of this complexity.

Measurements of transpiration and photosynthetic rates revealed species-specific responses to different light colors during the day, indicating a significant interaction between species and light color treatments (table 1). As a result of these differences, some plant species may be less affected by ALAN, making them better suitable for use in urban environments. These findings also indicate that ALAN can influence daytime photosynthesis, potentially via the downregulation of daytime photosynthetic activity (Park et al., 2021). However, no evidence was found to suggest that the photosystems of ALAN-exposed plants exhibited altered nighttime photosynthetic rates (Figure 2f). Instead, the observed patterns may be explained by an increase in stomatal density, a physiological response linked to adaptation to low-light environments (Shimazaki et al., 2007).

Previous research has recommended the use of streetlights with a significant red component since animals react least to red light (Durmus et al., 2024). However, our findings show that the use of red light can actually negatively impact plants, since this color had the highest impacts on plant morphology as well as as assimilation. Since plants rely on the entire light spectrum to regulate their growth, it is essential to carefully assess all colors (Diamantopoulou et al., 2021). Our study demonstrates that both light intensity and color significantly influence plant morphology and assimilation. While some treatments showed no significant effects, long-term effects may still be present. Detailed dose-response curves for low light intensities are needed to find a balance between functionality of light in cities, and ecological impacts. It is important to understand how the effects of ALAN can be mitigated by different colors or light intensities. As this study shows, even light conditions that appear harmless and are even promoted to mitigate ecological impacts, may have unexpected effects on plant . Therefore, we need to improve our understanding of ALAN’s effect on plants, a taxa that has been under-researched in this field yet is the foundation for ecosystem functioning, and to ensure this knowledge reaches decision-makers to protect wildlife within urban areas.

## Acknowledgements

This project was funded by the Bioclock Consortium through the NWA-ORC program of the Dutch Research Council (NWO; project number 1292.19.077). We would like to express our gratitude to Dr. Fredric Lens and Dr. Giovanni Bortolami for providing the equipment necessary for data collection and for their guidance on measurement techniques. We also extend our thanks to Dr. Kamiel Spoelstra for granting us access to the experimental sites, and to Zeyneb Gokce for her invaluable assistance during fieldwork.

## Author contributions(CRediT)

**E. Castrop**: Conceptualization, Data curation, Formal analysis, Investigation, Methodology, Visualization, Writing – original draft, Writing – review and editing. **P. M. van Bodegom**: Funding acquisition, Formal analysis, Writing – original draft, Writing – review and editing, Supervision. **E. F. Strange**: Conceptualization, Writing – original draft, Writing – review and editing, Supervision. **S. H. Barmentlo**: Conceptualization, Data curation, Formal analysis, Supervision, Writing – original draft, Writing – review and editing

## Data availability

All data is available in the Dryad open data repository, https://doi.org/10.5061/dryad.cz8w9gjdz

